# A Single Gene Causes an Interspecific Difference in Pigmentation in *Drosophila*

**DOI:** 10.1101/014464

**Authors:** Yasir H. Ahmed-Braimah, Andrea L. Sweigart

**Author notes:** Corresponding author: Department of Biology, 340 Hutchinson Hall, University of Rochester, RC Box 270211, Rochester, NY 14627.

## Abstract

The genetic basis of species differences remains understudied. Studies in insects have contributed significantly to our understanding of morphological evolution. Pigmentation traits in particular have received a great deal of attention and several genes in the insect pigmentation pathway have been implicated in inter- and intraspecific differences. Nonetheless, much remains unknown about many of the genes in this pathway and their potential role in understudied taxa. Here we genetically analyze the puparium color difference between members of the Virilis group of *Drosophila*. The puparium of *Drosophila virilis* is black, while those of *D. americana, D. novamexicana*, and *D. lummei* are brown. We used a series of backcross hybrid populations between *D. americana* and *D. virilis* to map the genomic interval responsible for the difference between this species pair. First, we show that the pupal case color difference is caused by a single Mendelizing factor, which we ultimately map to an ~11kb region on chromosome 5. The mapped interval includes only the first exon and regulatory region(s) of the dopamine *N*-acetyltransferase gene (*Dat*). This gene encodes an enzyme that is known to play a part in the insect pigmentation pathway. Second, we show that this gene is highly expressed at the onset of pupation in light-brown taxa (*D. americana* and *D. novamexicana*) relative to *D. virilis*, but not in the dark-brown *D. lummei*. Finally, we examine the role of *Dat* in *adult* pigmentation between *D. americana* (heavily melanized) and *D. novamexicana* (lightly melanized) and find no discernible effect of this gene in adults. Our results demonstrate that a single gene is entirely or almost entirely responsible for a morphological difference between species.

## Introduction

Understanding the genetic basis of morphological differences between species is a central goal of evolutionary biology. A key question focuses on complexity: do species differ by many genes of small effect or by a few genes of large effect (Haldane 1937)? The answer to this question has remained elusive; cases studied present a mixture of both scenarios (Orr 2001). This is not surprising, however, as traits and species may differ in divergence rates and types of selection (or lack thereof). The question can best be stated as one of relative frequency: how often do major genes cause morphological differences between species (Orr 2001)?

Insect pigmentation traits have received much attention from geneticists (Wittkopp *et al*. 2003a; True 2003; Wittkopp and Beldade 2009). These studies benefited from the fact that the pathway that determines insect cuticular pigment is both conserved and well understood (Wittkopp *et al*. 2002; Wittkopp *et al*. 2003; Jeong *et al*. 2008; Williams *et al*. 2008; Wittkopp *et al*. 2009; Werner *et al*. 2010). Several genes in the pigmentation pathway have been implicated in pigmentation differences between and within species, particularly in *Drosophila* (Wittkopp *et al*. 2003b; Jeong *et al*. 2008; Takahashi et al. 2007; Telonis-Scott *et al*. 2011; Bastide *et al*. 2013). Furthermore, in cases where the molecular basis of a phenotypic difference has been determined, the results have usually shown that regulatory mutations cause differences in expression between the alternative alleles (Wittkopp *et al*. 2003b; Pool and Aquadro 2007; Jeong et al. 2008; Werner et al. 2010; Takahashi and Takano-Shimizu 2011; Arnoult et al. 2013). As the pathway is currently understood, the interplay between a handful of enzymes (e.g. Ebony, Yellow, Tan), which act on tyrsoine and its derivatives such as dopa and dopamine, and a set of regulatory proteins (e.g. Bab) determine the amount and spatial distribution of cuticular pigment (Wittkopp *et al*. 2002; Wittkopp *et al*. 2003a). Variation in color pattern and pigment observed among insects, therefore, likely reflects evolutionary changes in the genes belonging to this pathway (Wittkopp, *et al*. 2003a).

The Virilis group of *Drosophila* provides a good model for analyzing the genetic basis of morphological differences among closely related species (Spicer 1991; Wittkopp *et al*. 2010; Fonseca *et al*. 2013). The Virilis group contains two phylads, the Montana phylad and the Virilis phylad. The Virilis phylad – the focus of this study – includes *D*. virilis, *D. lummei*, *D. novamexicana* and *D. americana*. Several studies have examined the phylogenetic history and reproductive incompatibilities among members of this group (Orr and Coyne 1989; Caletka and McAllister 2004; Sweigart 2010; Morales-Hojas *et al*. 2011; Sagga and Civetta 2011; Ahmed-Braimah and McAllister 2012). The sequenced genome of *D. virilis*, coupled with the relative ease of producing backcross and advanced generation hybrids in species crosses, allows for easy genetic marker development and mapping of genomic regions underlying trait differences. This group, however, contains many segregating and fixed chromosomal inversions, which sometimes render fine-scale genetic mapping in parts of the genome difficult or impossible (Hsu 1952).

Species belonging to the Virilis phylad differ in several morphological characters (Patterson *et al*. 1940; Spencer 1940; Stalker 1942). Perhaps best known, adults of *D. novamexicana* differ in color from their sister species. While all other members of the group are darkly pigmented, *D. novamexciana* has evolved a lightbrown color along its dorsal abdomen, head and thorax. *D. novamexicana* also lacks pigment along the abdominal dorsal midline (Spicer 1991; Wittkopp *et al*. 2003). Previous work has shown that *ebony* plays a major role in this pigment difference (Wittkopp *et al*. 2003), and that *tan* also contributes substantially (Wittkopp et al. 2009).

Species belonging to the Virilis phylad also differ in pupal case color (see Figure S1 & 5a)(Stalker 1942). All members of the group (including Montana phylad species) have brown puparia. *D. virilis*, on the other hand, lacks brown pigment and has black puparia that are easily distinguished from the sister species early in pupal development. Although this trait appears fixed in *D. virilis*, the intensity of the black color varies between strains and is also affected by crowding and/or culture conditions (Spencer 1940; Stalker 1942). For example, uncrowded pupae appear darker than crowded ones, which appear tannish gray.

Previous genetic analyses of the pupal color difference between *D. americana* and *D. virilis* conducted during the early 1940s yielded mixed results. Early work by W. P. Spencer suggested a complex genetic basis of this trait, while his later work did not support this conclusion (Spencer 1940). Spencer attributed the discrepancies in his findings to the possibility that pupal color may behave differently in different strains. Interestingly, Spencer’s original findings on this trait played an important role in shaping H. J. Muller’s “multi-genic” view of morphological evolution (Muller 1940; Orr and Coyne 1992). Around the time that Spencer conducted his experiments, H. D. Stalker performed an independent genetic analysis of the pupal color trait between *D. americana* and *D. virilis* and attributed the difference to a large effect on chromosome 5 (Muller Element C) with supplementary genes for “brownness” carried on the *D. americana* chromosome 2-3 (Muller Elements D-E)(Stalker 1942). His results differed from those of Patterson, Stone and Griffen, who suggested that the difference is entirely explained by chromosomes 2-3(Patterson *et al*. 1940). The latter authors based their conclusions on cytological data, but provided no detailed account of their findings.

Here we present a genetic analysis of the pupal case color difference between two species of the virilis group, *D. americana* and *D. virilis*. We first map this difference at the level of whole chromosomes and find that only chromosome 5 causes the difference. We ultimately identify a single genomic interval (~11kb) on chromosome 5 that causes the pupal case color difference. This region contains only the first exon and regulatory region of a single gene, *GJ20215*. *GJ20215* is the homologue of the *D. melanogaster* dopamine *N*-acetyltransferase (*Dat*). *Dat* (also known as arylalkilamine N-acetyltransferase, or *aaNAT*) is an enzyme known to act within the pigmentation pathway. In particular, *Dat* catalyzes the reaction from dopamine (DA) to N-acetyl dopamine (NADA). We then examine the expression differences between pupae of *D. americana* and *D. virilis*, their hybrids, as well as among members of the virilis phylad for *Dat*. Finally, we test the role of this gene in the adult pigmentation difference between *D. americana* and *D. novamexicana*.

We conclude that reduced expression of *Dat* in early pupal development in *D. virilis* is the cause of dark pupae in this species.

## Materials and Methods

### Fly strains and husbandry

Flies were maintained at a constant temperature (22°C) in a ~12h day/night cycle on standard cornmeal medium. The *D. virilis* strain used (15010-1051.31) was obtained from the UCSD Drosophila Species Stock Center (http://stockcenter.ucsd.edu) and carries two visible mutations on the fifth chromosome: Branched (*B*) and Scarlett (*st*). Both mutations behave recessively in a *D. americana* genetic background, although *B* is considered dominant within *D. virilis* strains. (*B* causes abnormal branching of wing veins, and *st* mutants have bright red eyes.) The *D. americana* strain (SB02.06) was collected by Dr. Bryant M. McAllister (University of Iowa) in 2002 near the Cedar River in Muscatine County, Iowa. While *D. americana* harbors a number of chromosomal inversions and two chromosomal fusions, the strain used here differs from *D. virilis* in a small inversion on the telomeric end of chromosome 5 (In5a) and a large inversion encompassing the centromeric half of chromosome 2. Importantly it lacks the large In5b inversion that segregates in western populations of *D. americana* and affects~60% of the chromosome. In addition to the fixed fusion of chromosomes 2 and 3, the strain is also fixed for the clinally-distributed fusion between chromosomes 4 and X, both of which carry fixed inversion differences (see Figure 1a for karyotypes). The *D. novamexicana* (15010-1031.04) and *D. lummei* (LM.08) strains were obtained from Dr. Bryant M. McAllister. For all crosses in this study, male and female flies were collected within 2 days of eclosion and reared separately until they reached sexual maturity and crossed at 12-14 days old.

### Molecular genotyping

Up to 42 microsatellite markers and 6 single nucleotide polymorphism (SNP) markers were used to genotype recombinant individuals along chromosome 5(Table S1; see Figure S3 for arrangement of genome scaffolds on chromosome 5). Tandem repeat regions were identified in the *D. virilis* genome (Release droVir2, UCSC) using Tandem Repeat Finder (Benson 1999), and primers flanking repeat regions were designed using Primer3 (Rozen and Skaletsky 2000). SNPs were identified by sequencing small orthologous fragments of *D. virilis* and *D. americana* and identifying base substitutions or insertions/deletions unique to either strain used in the study. Genomic DNA was obtained from each individual whole fly using the protocol of Gloor & Engels (1992). All microsatellite markers were amplified using a touchdown PCR reaction in which annealing temperatures were incremented from 62°C to 52°C for 10 cycles and subsequently annealed for an additional 30 cycles at 52°C. Species-specific alleles were identified by sizing PCR-amplified fragments with an incorporated 5’ fluorescently labeled primer on an ABI-3700 automated capillary sequencer (Applied Biosystems). The genomic regions containing SNP and/or insertion/deletion markers were amplified using the NEB LonAmp-Taq protocol (ww.NEB.com), and fragments were purified (Qiagen PCR purification Kit) before they were sequenced on an ABI-3700 machine.

### Whole-chromosome association backcross

The backcross population of pupae segregating for whole-chromosomes was generated by backcrossing “AV” F1 males (A = *D. americana*, V = *D. virilis*; female genotype given first hereafter) to *D. virilis* females. As the *D. virilis* allele is recessive (Stalker 1942), this cross produces a population of pupae that are either black (homozygous for *D. virilis* at the pupal case color locus, n = 81) or light-brown (heterozygous at the pupal case color locus, n = 84). To infer the whole-chromosome genotypes of each individual, pupae were genotyped at a single microsatellite marker for each of the three autosomal linkage groups (chromosomes X-4:SSR21, chromosomes 2-3:SSR37, and chromosome 5:SSR146). Chromosome 6 (the small “dot” chromosome, Muller Element F) was not surveyed.

### Recombinant mapping of the pupal case color locus on chromosome 5

A recombinant backcross mapping population between *D. americana* and *D. virilis* was generated for another purpose, but was used in this study to genetically map the interval containing the pupal case color locus. Recombinant backcross males ((AV)A) were generated by backcrossing AV F1 females to *D. americana* males. As all backcross progeny possessed the light-brown pupal case color (caused by the dominant *D. americana* allele), the pupal case color alleles inherited by (AV)A males can only be assessed by crossing recombinant backcross males to *D. virilis* females; (AV)A males heterozygous at the pupal case color locus will sire light-brown and black pupae in equal proportions whereas males homozygous for *D. americana* alleles will produce only light-brown pupae. A total of 644 recombinant backcross males ((AV)A) that sired progeny comprised the entire mapping population, however only a subset of individuals was genotyped along the majority of the recombining segment of the chromosome. This subset comprised 46 individuals and was selected based on the genotype at the visible marker, *B*, and whether that individual sired ≥ 5 progeny (only *D. americana* homozygotes at *B* were selected). Only males that sired progeny were selected for different reasons (see(Sweigart 2010)), but also because progeny were used to identify the paternal pupal case genotype. In addition, a second subset of 24 individuals was selected based on known recombination events in the vicinity of the pupal case locus from past genotyping efforts in the laboratory. The 70 individuals were genotyped at up to 44 microsatellite and single nucleotide polymorphism (SNP) markers along a 14.5Mb region that encompasses ~70% of the recombining portion of the chromosome.

### Screening for recombinants in the candidate region

To screen for additional recombinants in the 27kb candidate region, a large collection (n ~ 30,000) of recombinant backcross pupae was generated by backcrossing F1 AV females to *D. virilis* males. Pupae were separated by color (light-brown or black) and allowed to eclose. The phenotype of eclosing flies at the two visible markers (*st* and *B*) was recorded and only flies that were recombinants between *st* and *B* (~30cM) were frozen for molecular genotyping (n ≈ 12,000). Recombinants between *st* and *B* were genotyped at two microsatellite markers (SSR169 and SSR173) which are ~32kb apart and flank the pupal case color locus. A total of 16 recombinants was recovered between SSR169 and SSR173, and were subsequently genotyped at 10 additional markers (6 SNP, 4 microsatellite) spanning the 32kb region.

### Multiple sequence alignment of candidate region in *D. americana* and *D. virilis* strains

Publicly available genome sequences for two *D. americana* strains (H5 and W11, http://cracs.fc.up.pt/~nf/dame/fasta/; (Fonseca et al. 2013)) and two *D. virilis* strains (Str9: SRX496597, and Str160:SRX496709; (Blumenstiel 2014)) were used to compare the candidate region between the two species. To extract orthologous regions in the *D. americana* strains, the sequence of the ~11kb candidate region from the *D. virilis* genome strain (droVir2) was Blasted (BLAST 2.2.28+) against the H5 and W11 genome assemblies. Matching scaffolds for each *D. americana* strain were concatenated and trimmed accordingly (Geneious R8). Sequence reads for the two *D. virilis* strains were obtained from the Sequence Read Archives (SRA) and mapped to the *D. virilis* genome using bwa (v.0.7.9a). The consensus sequence from the mapped reads of the ~11kb candidate region for each *D. virilis* strain was obtained through “pileup” of consensus bases and converted to FASTA sequences using Samtools (v.0.1.19). Finally, the ~11kb region from all four strains and the *D. virilis* genome strain were aligned using MUSCLE (v.3.8.31).

### Quantitave RT-PCR

The orthologue of *GJ20215* in *D. melanogaster* is known to have two isoforms, yet the annotated *GJ20215* consisted of only one isoform (hereafter isoform-A, FlyBase v.2014_04). To determine the gene structure of *GJ20215*, publicly available *D. virilis* mRNA short sequence reads were obtained from the modENCODE project (www.modencode.org). Reads were processed and mapped to the *D. virilis* genome using Tophat (Trapnell *et al*. 2009) and transcripts assembled using Cufflinks (Trapnell et al. 2010). The gene structure of *GJ20215* was found to contain a second isoform (isoform-B) that utilizes an alternative, previously unknown first exon which lies ~12.5kb upstream of the first exon of isoform-A. Two different isoform-specific forward primers that overlap the exon-exon boundary, in addition to a shared reverse primer, were designed as described above. These primers amplify a ~150bp mRNA fragment. Primers were also designed for a reference gene, *RpS3*. Amplification efficiencies were estimated for each primer pair using the standard curve method. Total RNA was extracted from each virilis group strain (~10 individual pupa/replicate) at three different stages during pupal development:(1) at the onset of cuticle hardening but prior to the development of any pigment, (2) after the development of light pigment (~6-8 hrs after cuticle hardening), and (3) after full pigment development (~12-16 hrs after cuticle hardening). In addition to pure species samples, total RNA was also extracted from F1 hybrid pupae (all three stages) and F3 intercross hybrid pupae (only stage “3”) between *D. americana* and *D. virilis*. For each extraction pupae were flash-frozen in liquid nitrogen and RNA extracted using the Qiagen RNeasy Mini kit (Qiagen). Approximately 4ug of RNA from each sample was used to synthesize cDNA that was used in SYBR-green quantitative RT-PCR (qRT-PCR) reactions (Qiagen). Samples were grouped accordingly for each comparison with 2-5 replicates run in a single plate in two technical duplication runs. Ct values for each replicate were averaged across biological replicates and technical replicates. Relative quantitation was performed using the ∆∆Ct method (Livak and Schmittgen 2001) with RpS3 as the reference gene and *D. americana* as the control sample. Statistically significant differences in relative expression were assessed by performing a two-sample t-test (R v.3.1.1).

### *D. americana/D. novamexicana* adult pigmentation analysis and genotyping

Sixth generation (F6) intercross hybrids between *D. americana* and *D. novamexicana* were produced by crossing *D. americana* females to *D. novamexicana* males en masse and subsequently allow hybrids to intercross for five generations. F6 individuals were separated by sex soon after eclosion and aged for 14 days. 188 flies (93 females, 95 males) were imaged dorsally before flash-frozen for DNA extraction, and red-blue-green (RBG) measurements were obtained (ImajeJ) from three landmarks: the scutellum, the anterior portion of the throax, and the dorsal abdominal cuticle. Only red measurements were used in the analyses as they showed the greatest difference. A microsatellite marker (for *tan*) and two sequenced fragments containing multiple SNP differences (for *ebony* and *GJ20215*) were developed to genotype F6 individuals as described above. The data were analyzed by performing a one-way analysis of variance (ANOVA) for each of the three markers separately where the genotype at a given marker is the independent variable and red mean value as the dependent variable (R.v.3.1.1).

## Results

### Mapping to whole chromosomes

The *D. virilis* pupal case color allele(s) is completely recessive(Spencer 1940; Stalker 1942). Reciprocal F1 hybrid pupae between the two species are identical in pupal color to *D. americana*. In addition, backcross hybrids are either black (*D. virilis* homozygotes at pupal color locus) or brown (heterozygotes), with no observable gradation between these two classes. These observations suggest that the factor(s) causing the difference in pupal color in this species pair reside(s) on autosomes.

To map this phenotypic difference to whole autosomes, we generated a backcross population of pupae by crossing F1 AV hybrid males to *D. virilis* females (Figure 1a). As meiotic recombination does not occur in *Drosophila* males, whole chromosomes remain intact and species-origin can be identified with single markers per chromosome.

Chromosome 5 (Muller Element C) is perfectly associated with pupal case color (Figure 1b). Chromosomes 2-3 and 4 (Muller Elements E-D and B), on the other hand, show no association with the phenotype. We also observe an excess of heterozygous genotypes for chromosome 2-3, but this excess is not correlated with segregation of black and light-brown pupae. The gene(s) that cause the pupal color difference between species thus reside on a single autosome: chromosome 5.

### Fine-mapping the pupal case color locus

*D. americana* harbors a number of fixed and segregating chromosomal inversions relative to *D. virilis* (Hsu 1952). Such differences can often preclude finer recombination mapping. Fortunately, the *D. americana* strain used in this study is homosequential with *D. virilis* along ~85% of the euchromatic arm of chromosome 5. We can therefore fine-map the locus (or loci) underlying the pupal color difference between *D. americana* and *D. virilis*. To do so, we performed two rounds of recombination mapping using two different recombinant backcross populations. We describe each below.

In the first round we used a recombinant backcross population (generated for different purposes) that was created by backcrossing F1 females (AV) to *D. americana* males (Figure 2a). Because *Drosophila* females *do* recombine, backcross hybrids can carry recombinant chromosomes. Backcross hybrids were either homozygous for *D. americana* alleles or heterozygous at any given locus, and therefore all had brown pupae. To infer their genotype at the putative pupal color locus we examined the pupal colors of the progeny sired when individual backcross males were crossed to *D. virilis* females. Backcross males that were heterozygous at the pupal color locus sired both black and brown pupae, while backcross males homozygous at the pupal color locus sired only brown pupae.

The *D. virilis* strain used in this study carries two visible markers on chromosome 5, *Branched* (*B*) and *Scarlett* (*st*), both of which behave recessively in a *D. americana* genetic background. *B* resides near the center of the chromosome (Figure 2b). We selected individuals from the recombinant backcross male population that were homozygous for *D. americana* alleles at the *B* locus (n = 46 such males), but carried both possible genotypes at the pupal color locus. These individuals were genotyped using microsatellite markers that spanned most of the length of the chromosome. This analysis revealed that the pupal color locus resides in a 2.5Mb region nearly midway between *B* and *st* (Figure S2). To refine our mapping here further, we studied an additional set of backcross males that was sparsely genotyped along chromosome 5 but carried a recombination breakpoint within the 2.5Mb region (n =24). The entire set of genotyped individuals is represented in Figure 2b. This mapping population lets us narrow the pupal case color locus to a genomic region that spans 27kb (Figure 2c). This region includes 4 genes: *GJ22136*, *GJ22137*, *GJ20214* and *GJ20215* (*Dat*).

In the second round of mapping we generated another recombinant mapping population by backcrossing F1 AV females to *D. virilis* males and separated pupae based on color (Figure 3a). Performing the backcross in this way allows us to recover both color phenotypes among backcross individuals. Furthermore, we enriched for recombinants in the relevant region by only genotyping eclosed adults that had a recombination event between our two visible markers. These individuals were then genotyped at two microsatellite markers that flank the 27kb candidate interval. Out of ~30,000 pupae, we recovered 16 recombinants between the two microsatellite markers. The 16 recombinant individuals were subsequently genotyped along the 27kb candidate region (Figure 3b). Using this approach we narrowed the location of the relevant locus to an ~11kb region.

This region contains only the first intron, first exon and upstream regulatory region of the annotated *Dat* isoform (isoform-A). This region also includes the majority of the first intron of the unannotated isoform of this gene (isoform-B). Divergence in pupal case color between *D. virilis* and *D. americana* is thus caused by sequence differences in this ~11kb region.

### Sequence alignment of mapped interval between and within *D. americana* and *D. virilis*

The mapped interval implicates *Dat* as causal and excludes adjacent genes. This interval mostly contains sequence that is noncoding, but also includes the first exon of isoform-A of *Dat*. To examine the extent of sequence divergence between the two species in this interval, we generated a multiple sequence alignment of the region using publicly available genome sequences from two strains of *D. americana* (strain H5 and strain W11; (Fonseca et al. 2013)) and two strains of *D. virilis* (Strain-9 and Strain-160; (Blumenstiel 2014)). By Blasting the candidate region’s sequence from the *D. virilis* genome to the two *D. americana* genomes, we recover two and three scaffolds from the H5 and W11 strains, respectively, that cover the ~11kb region in *D. virilis*. The two H5 scaffolds (H5C.563 and H5C.564) cover all but 62bp of the candidate region, whereas the three W11 scaffolds (W11C.1012, W11C.1013, and W11C.1014) cover all but ~200bp. *D. virilis* Strain-9 and Strain-160 reads covered the entire candidate region.

We found a total of 1800 sequence differences in the candidate region between and within the strains sampled here (Figure 4a, File S1). We classified base substitutions and insertion/deletions separately, and partitioned them into fixed mutations, polymorphic in *D. americana*, polymorphic in *D. virilis*, and shared polymorphisms. The largest subset was nucleotide substitutions fixed between *D. americana* and *D. virilis* (n=469; Figure 4a), while *D. americana* showed more within-species polymorphism than did *D. virilis*. The number of shared polymorphisms was also low (n=81) but similar to polymorphism levels within *D. virilis*.

The sequence alignment reveals two nucleotide substitutions that are fixed between *D. americana* and *D. virilis* in the open reading frame of the first exon (Figure 4b). One of these mutations causes a nonsynonymous substitution (His -> Arg) at the 7^th^ codon, while the other is synonymous. Mutations elsewhere in the mapped interval are distributed roughly uniformly throughout the 5’UTR of *Dat* and along the introns. Although the number of fixed nucleotide substitutions is nearly twice that of insertions/deletions, the latter affects ~60% of aligned variable sites (Figure 4b, File S1). These results show that many fixed differences exist between *D. americana* and *D. virilis* in the mapped interval, with all but one affecting noncoding sequences.

### Relative expression of *Dat*

To examine the possible role of gene regulation in causing the two pupal color phenotypes, we measured relative expression levels of *Dat* using qRT-PCR at three phenotypically defined stages in *D. americana* and *D. virilis*, their hybrids, and in other species that belong to the Virilis phylad. We defined first stage pupae (“pre-pupae”; *pre*) as those with hardened cuticles but no visible pigment, second stage pupae (“pupae”; *pup*) as slightly pigmented (faint brown; both species appear identical), and third stage pupae as fully pigmented (“post-pupae”;*pos*) (Figure 5a).

First, we measured relative gene expression levels for both *Dat* isoforms in *D. americana* and *D. virilis* pupae. We found that isoform-A differs drastically in expression at all three stages (*p*<*0.001*) (Table S2). Specifically, *D. americana* pupae express *Dat* isoform-A >80-fold higher than *D. virilis* in all three stages. Expression of Isoform-B, on the other hand, differs modestly but significantly only in the first and third stages (*p*=*0.0016* and *p*=*0.017*, respectively; Figure 5b, Table S2). Isoform-B is on average ~1.5-fold and ~3-fold higher in *D. americana* at the first and third stages, respectively. These results suggest that isoform-A is likely the isoform affected most by regulatory sequence divergence between *D. americana* and *D. virilis*.

Second, we examined expression levels of isoform-A among F1 hybrids in all three stages of pupal development. We also examined expression levels in brown F3 hybrids and black F3 hybrids, but only in the third stage as this is the stage at which pupal color can be determined (Figure 5c). F1 hybrid pupae fully resemble pure *D. americana* pupae. Likewise, their expression of *Dat* isoform-A did not significantly differ from that of *D. americana* in the first and second stage (*P*>*0.05*, Table S2). However, F1 hybrid expression is reduced markedly at the third stage (*p*=*0.08*). Similar to F1 hybrids, brown F3 hybrid pupae had markedly reduced expression of *Dat* isoform-A at the third stage relative to *D. americana* (*p*=*0.06*). In contrast, black F3 hybrid pupae had expression levels similar to pure *D. virilis* at the third stage (*p*=*0.07*). These results suggest that expression levels at the third stage do not account for most of the phenotypic difference between black and brown pupae. Instead, expression levels earlier in pupal development likely determine this phenotypic difference.

As noted, the black pupal case color of *D. virilis* is unique in the Virilis group: the sister species all feature brown pigment in their pupal cuticles. *D. novamexicana* pupae are indistinguishable from *D. americana. D. lummei* pupae, on the other hand, appear intermediate between *D. virilis* and *D. americana/D. novamexicana*,with pupae containing both brown and black pigment (Figure 5a). We examined the expression of *Dat* isoform-A in pupal samples from the three stages in all four virilis phylad species (Figure 5d). We found that expression levels in *D. novamexicana* samples were slightly higher than *D. americana* in the first two stages (*p*~0.1), but are slightly reduced in the third stage (*p*=0.1). *D. lummei* expression levels, on the other hand, closely resemble *D. virilis* samples and are uniformly low across all three stages (*p*>*0.05*; Figure 5d). This suggests that the pupal color difference between *D. lummei* and *D. virilis* may have a genetic basis that differs from that identified here.

Taken together these observations are consistent with the brown pupal case color resulting from increased expression of *Dat* (isoform-A) early in pupal development, whereas production of black pigment results from low expression.

### The role of *Dat* in adult pigmentation

Previous work on the genetic basis of the difference in *adult* pigmentation between *D. americana* and *D. novamexicana* showed that it is largely caused by two genes, *ebony* and *tan*. Indeed these two genes together account for nearly 80% of the pigmentation difference (Wittkopp et al. 2003b; Wittkopp et al. 2009). In the initial mapping study performed by Wittkopp *et al*. (2003b), a QTL region containing the *Ebony* locus showed the highest LOD score, with both *D. americana* and *D. novamexicana* alleles contributing to the abdominal pigmentation difference. Additional QTLs were observed on chromosomes 3 and 5. No pigmentation candidate genes were known on these chromosomes. Given *Dat*’s mode of action described here and elsewhere (see Discussion), we hypothesized that it might play a role in the color difference between *D. americana* and *D. novamexicana* adults, contributing to the remaining ~20% difference between the two species.

We quantified the difference in pigmentation between adults of *D. americana* and *D. novamexicana* by measuring the amount of emitted red light from three landmarks: the scutellum, the lightly pigmented portion of the thorax, and the abdomen (Figure 6). Pure species are clearly distinguishable in mean red values at all three landmarks, but the species differences are largest for abdominal coloration (Figure 6). F1 hybrids tend to have intermediate mean red values, though with larger variance.

We tested the association between the mean red value at the three adult landmarks and genotype at three markers tightly linked to *ebony*, *tan*, and *Dat* in F6 intercross hybrids (n = 188, Table S3). The mean red value for genotypes at *ebony* increased at all three landmarks as *D. novamexicana* alleles are added, but this effect was significant only for the abdomen (*p*=*0.002*; Figure 7; Table S3). A modest increase in mean red value was also observed with the addition of *D. novamexicana* alleles at *tan* (*p*=*0.003*; Figure 7; Table S3). In contrast, *Dat* showed no significant association with red values at any landmark (*p*>*0.55*; Figure 7; Table S3).

We conclude that *Dat* plays no discernable role in the pigmentation difference between *adults* of *D. americana* and *D. novamexicana*.

## Discussion

We have shown that a single gene entirely or almost entirely causes a morphological difference between two species of *Drosophila*. In particular, the difference between the black pupal case color of *D. virilis* and the brown pupal case color of *D. americana* maps to an ~11kb region that includes only a single exon and part of the regulatory region of the gene, *GJ20215*. *GJ20215* is the homologue of the *D. melanogaster* dopamine *N*-acetyltransferase gene (*Dat*). *Dat* is highly conserved among insects and is known to play a role in the insect pigmentation pathway where it catalyzes the conversion of DA to NADA (Wittkopp *et al*. 2003a; Wittkopp and Beldade 2009).

The mapped interval contains many fixed differences between *D. americana* and *D. virilis*. Among all fixed differences between the two species (710 total), only one causes an amino acid substitution; this change resides in the first exon. The first exon also contains a silent substitution. All other fixed differences reside within introns, the 5’UTR and the putative regulatory region(s). We have not identified the causal mutation(s) that explain the species difference. However, given the disproportionately large number of differences residing in noncoding regions, we examined differences in gene expression between the species.

The expression analysis shows that both isoforms of *Dat* show reduced abundance in *D. virilis* relative to *D. americana*. Across all stages surveyed, however, isoform-A shows a more dramatic expression difference between the two species. *Dat*’s apparent mode of action in the species pair studied here resembles two recent findings in other insects. First, recent work found that loss of the *Dat* orthologue in the silkworm, *Bombyx Mori*, results in increased production of black pigment in larvae and adults.(Dai *et al*. 2010; Zhan *et al*. 2010). Second, a recently developed dominant visible marker in insects uses a transgenic over-expression vector that expresses the *B. mori* orthologue of *Dat* (Osanai-Futahashi *et al*. 2012). This over-expression successfully reduces the production of black melanin in three distantly related species (*B. mori*, *D. melanogaster*, and *Harmonia axyridis*).

These studies suggest that *Dat* has a conserved phenotypic effect in a wide range of insects. Like other genes in the pigmentation pathway, *Dat* may sometimes behave as a “large-effect” gene, which – alone or with several other genes – can render closely related species morphologically different if sufficiently divergent. Several examples of large effect genes have been identified, mostly in *Drosophila* (e.g. Wittkopp *et al*. 2003b; Jeong *et al*. 2008; Werner *et al*. 2010), and often involve pigmentation phenotypes.

The adaptive significance of the pupal case color, if any, is not known. Pigmentation differences in adult insects have been attributed to a number of possible selective pressures, including desiccation resistance, ultra-violet protection, thermal regulation, crypsis and sexual selection (Hollocher *et al*. 2000; Brisson et al. 2005; Pool and Aquadro 2007; Wittkopp et al. 2010; Clusella-Trullas and Terblanche 2011; Telonis-Scott *et al*. 2011; Matute and Harris 2013). Some of these – such as desiccation resistance (Terblanche and Kleynhans 2009) and crypsis (Hazel *et al*. 1998) – may also apply to puparia. Some insect larvae/pupae are also targets of endoparasitoids and pupal case characteristics may affect infection propensity (Fellowes *et al*. 1999). Selection on pigmentation phenotypes may also be indirect as genes in the pigmentation pathway are often pleiotropic, affecting a number of traits (True 2003; Wittkopp and Beldade 2009). While some of the habitat preferences exhibited by virilis group members have been studied (Blight and Romano 1953), much remains to be understood regarding the ecological conditions that Virilis group species experience across all life stages.

In summary, our study shows that *Dat* is a potentially important player in pigmentation in insects. More important, our study also shows that the genetic basis of morphological differences between species is sometimes simple.

**Figure 1a:**
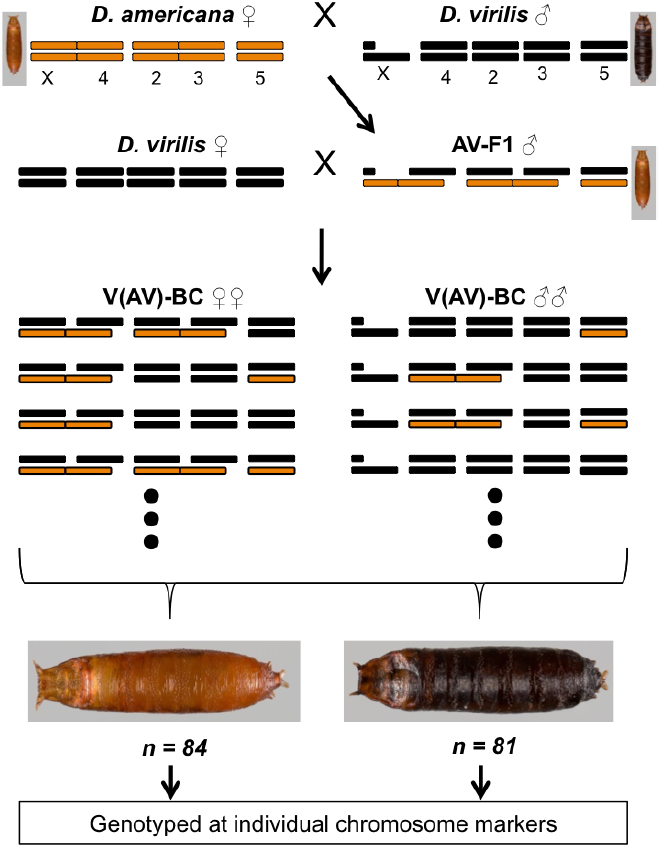
Crossing scheme to generate backcross pupae segregating for whole chromosomes: *D. americana* and *D. virilis* chromosomes are shown in light-brown and black, respectively. Female genotypes are shown on the left and male genotypes on the right. The pupal color phenotype for pure species and hybrids is shown next to the F1 karyotype. Eight possible genotypes (4 female, 4 male) are illustrated for backcross hybrids.

**Figure 1b:**
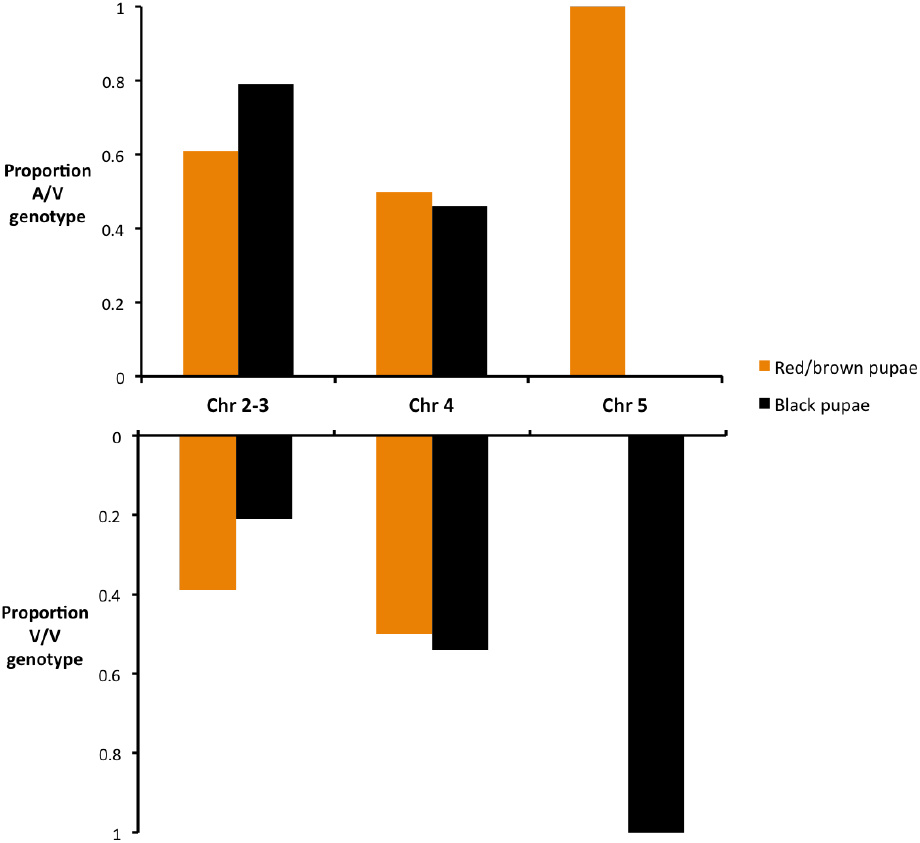
Associations between chromosomal genotypes and pupal color phenotype among whole-chromosome backcross hybrids: The proportion of the two alternative whole-chromosome genotypes for light-brown and black backcross pupae is plotted for each autosomal linkage group. The sum of A/V and V/V genotypes for each phenotypic class within each chromosome should be 1.

**Figure 2a:**
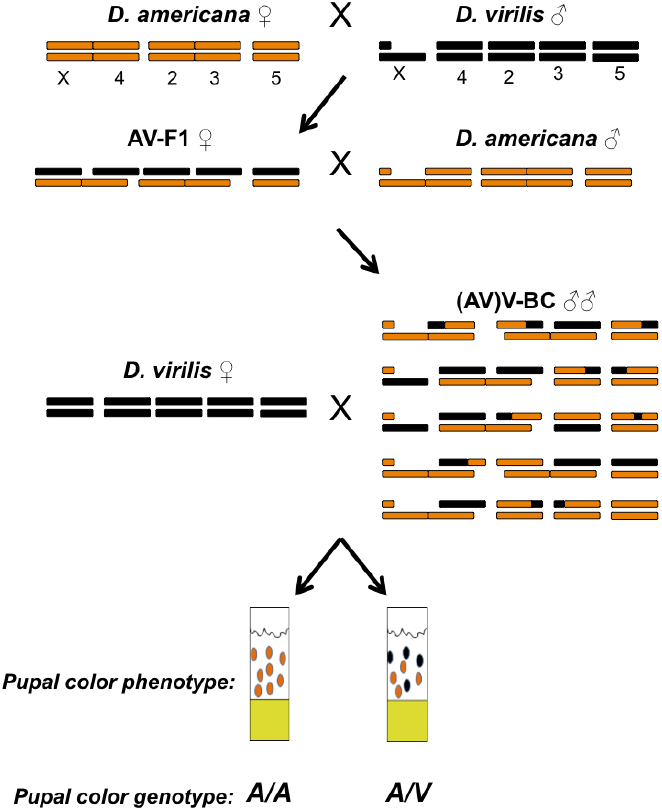
Crossing scheme to generate recombinant backcross hybrid males: Color scheme and positioning of male and female genotypes is the same as in Figure 1a. A sample of 5 possible recombinant backcross male genotypes is shown, and the test cross with *D. virilis* females is indicated. The two possible pupal color phenotypes and the corresponding genotypes at the pupal color locus are indicated at the bottom.

**Figure 2b, 2c:**
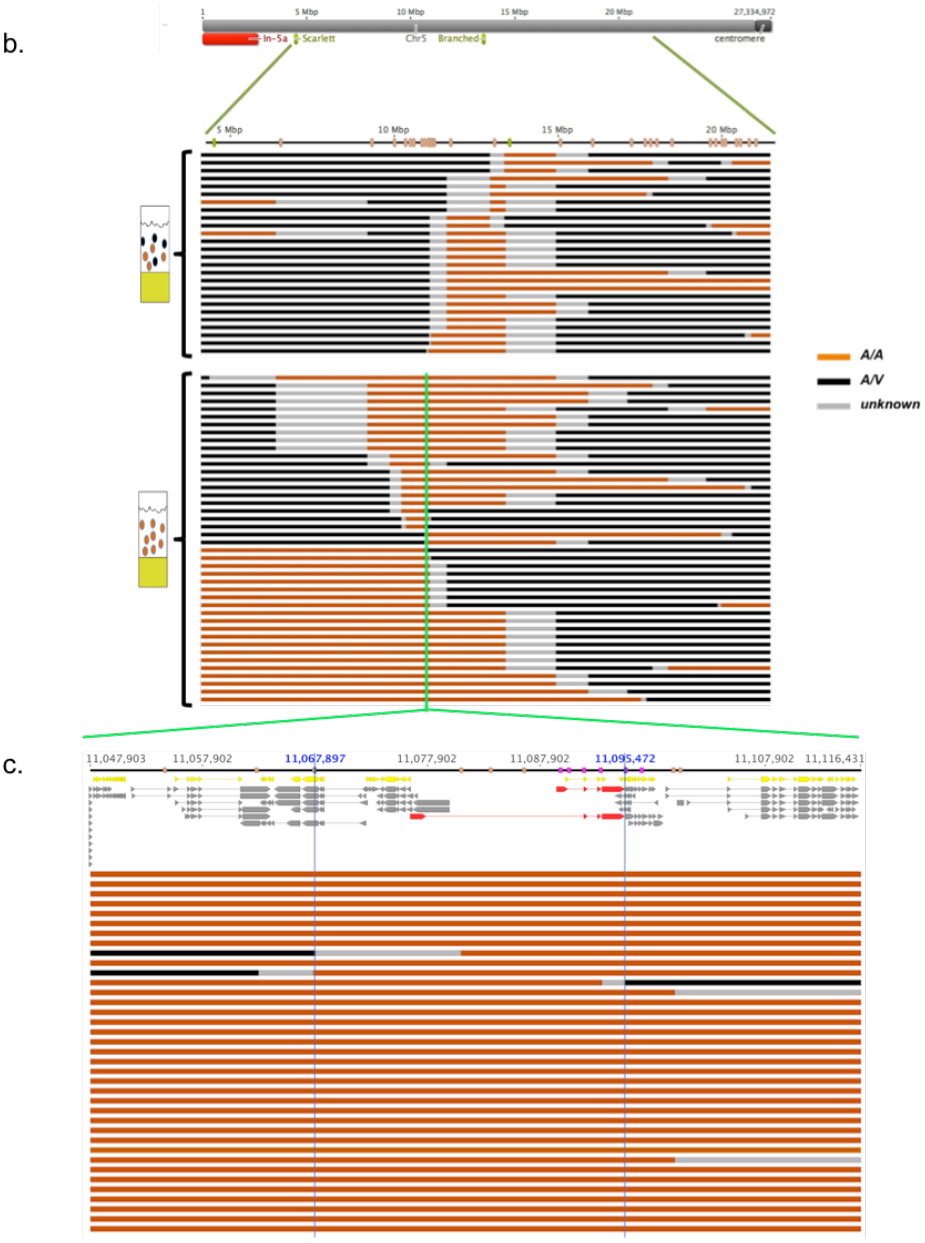
(b) Genotypes of recombinant backcross males along chromosome 5: The grey line at the top represents the entire length of chromosome 5, with the location of the two visible mutations indicated in green and inversion 5a indicated in red. Green lines magnify the molecularly genotyped region, where the location of microsatellite markers is indicated in beige. Each horizontal line below the chromosome represents a single recombinant backcross male, with regions along the chromosome color-coded according to inferred genotypes using microsatellite markers; light-brown = A/A, black = A/V, grey = unknown. Recombination breakpoints reside within grey regions. Recombinant males are grouped according to pupal color phenotype (shown on the left). The pupal color locus resides in a black region (A/V) among recombinant males who sire both black and light-brown pupae (upper group), or in a light-brown region (A/A) among recombinant males who sire lightbrown pupae only (lower group). (c) Close-up of a 68.5kb candidate region among recombinant backcross males who sired light-brown pupae only: The top portion shows the chromosomal coordinates and markers (beige=Microsattelite, purple = SNP) along with the gene annotations from Flybase (yellow) and those inferred by Cufflinks (grey). GJ20215 is shown in red (top = isoform-A, bottom = isoform-B).

**Figure 3a:**
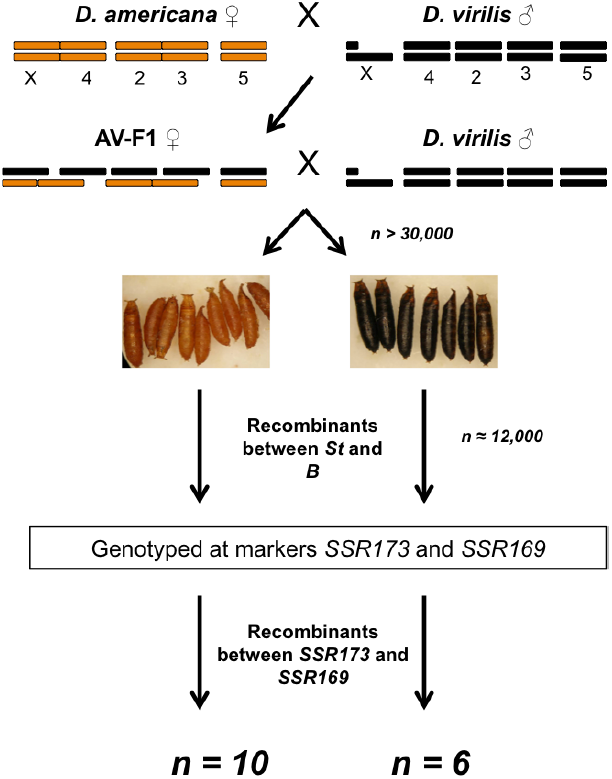
Crossing scheme to enrich for recombinants in the candidate region.

**Figure 3b:**
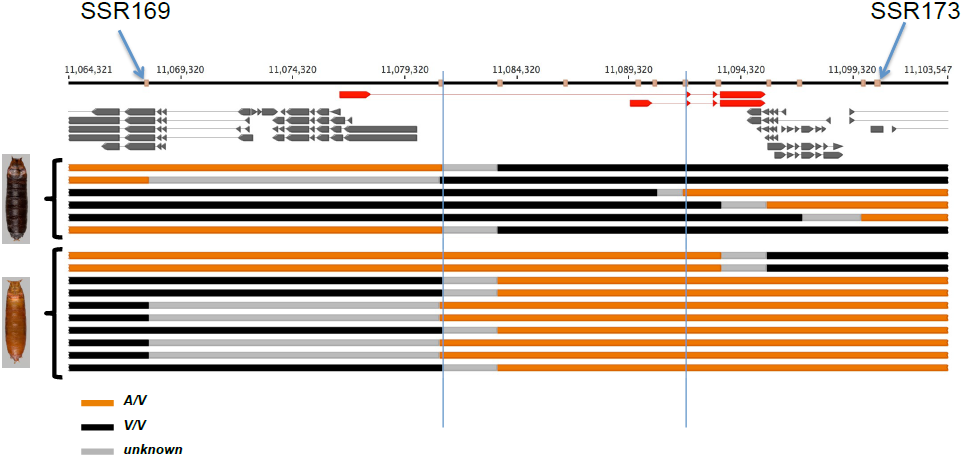
Recombinant individuals recovered using the recombination enrichment strategy: Only Cufflinks assembled transcripts are shown and GJ20215 is highlighted in red. The recombinants recovered are grouped by pupal color phenotype, where black indicates *D. virilis* homozygous regions (V/V) and light-brown are heterozygous regions. The two vertical blue lines indicate the mapped interval.

**Figure 4a:**
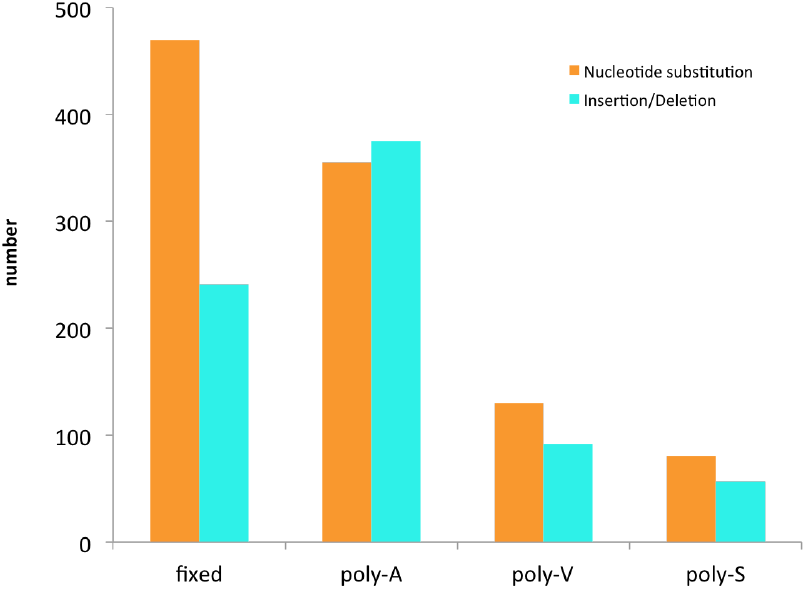
Total number of variable sites within the mapped interval across four classes: fixed differences, polymorphic in *D. americana* (poly-A), polymorphic in *D. virilis* (poly-V), and shared polymorphisms (poly-S). Variants are classified separately as nucleotide substitutions (orange) and insertion/deletions (blue).

**Figure 4b:**
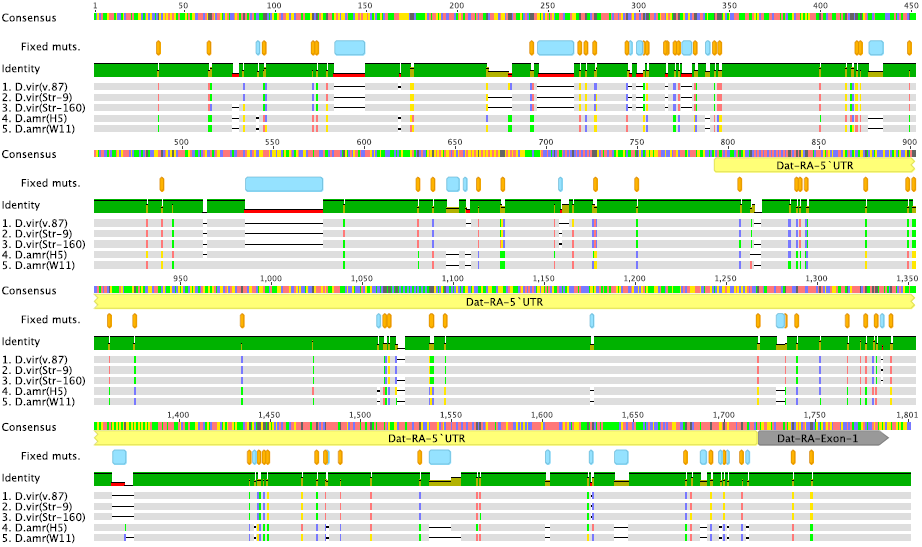
Multiple sequence alignment of a representative 1.8kb region: This alignment shows a portion of the ~11kb candidate region which includes ~800bp upstream of the transcription start site of *GJ20215* isoform-A, the 5’ untranslated region (yellow), and the first exon (grey). Fixed differences between *D. americana* and *D. virilis* are indicated in orange (nucleotide substitutions) and blue (insertion/deletions) (Geneious R8).

**Figure 5a:**
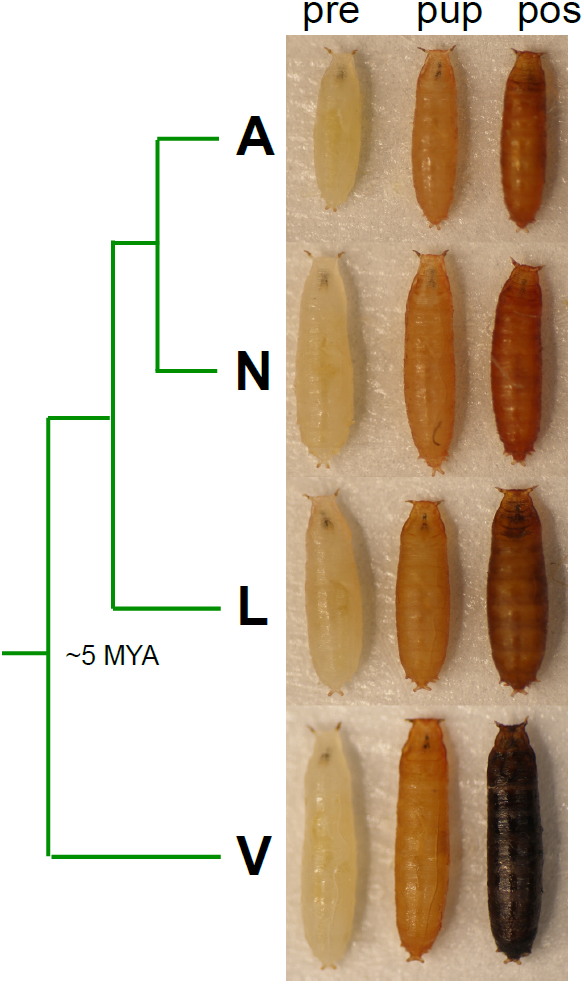
Pupae from the four virilis phylad members across the three developmentally defined stages used in RT-qPCR experiments (A = *D. americana*, N = *D. novamexicana*, L = *D. lummei*, V = *D. virilis*).

**Figure 5b, 5c, 5d:**
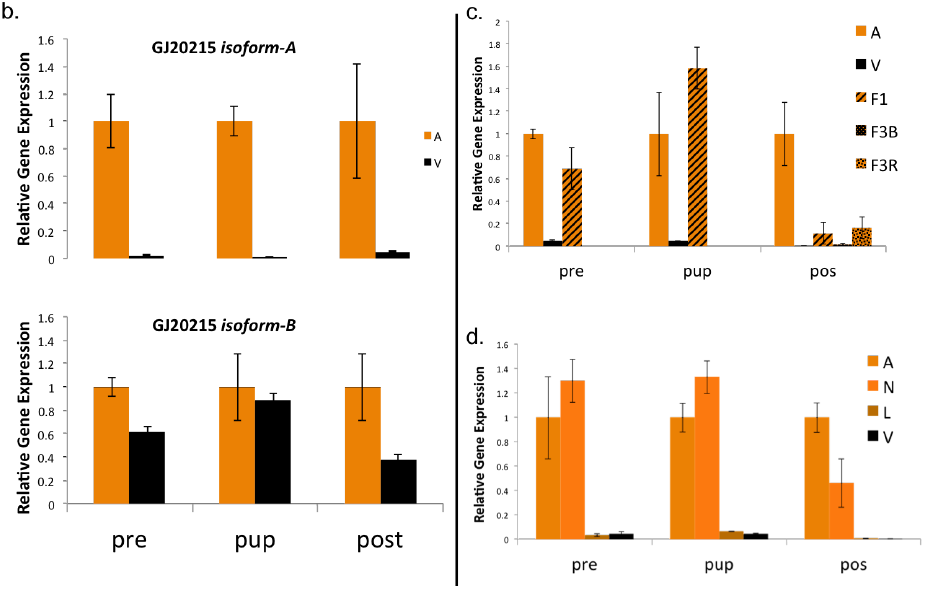
Relative gene expression measures for the two isoforms in *D. americana* (A) and *D. virilis* (V): Error bars represent standard error. All *D. americana* samples are normalized to 1 and considered the control sample. All comparisons are significant except for the “pup” sample of isoform-B (*p*<*0.05*). (c) Relative gene expression measures of *GJ20215* isoform-A in *D. americana*, *D. virilis*, first generation hybrids (F1), third generation hybrids with black pupae (F3B), and third generation hybrids with light-brown pupae (F3R): F3B and F3R samples were only assayed in the “pos” sample. Error bars and normalized controls are as in (b). (c) Relative gene expression measures in all four species of the virilis group (L = *D. lummei*, N = *D. novamexicana*).

**Figure 6:**
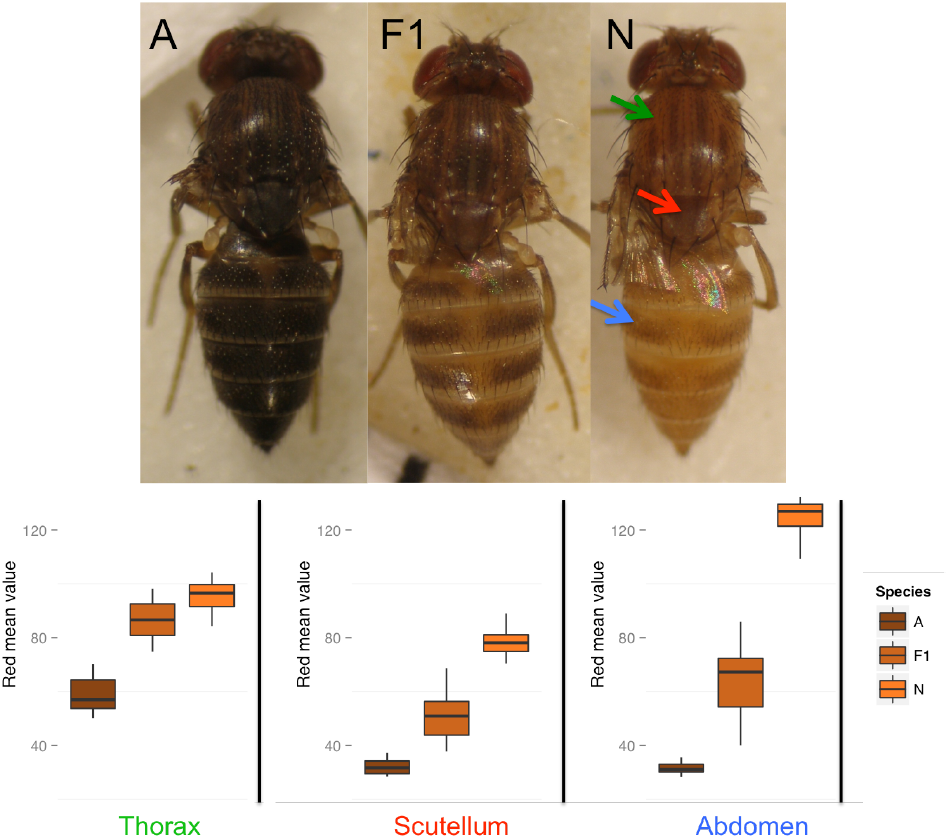
Distribution of mean read values across pure species and F1s: Top panel shows 14day old females of *D. americana* (A), *D. novamexicana* (N), and F1 hybrids (F1). Arrows in the “N” panel point to the adult landmarks used in quantifying emitted red values (arrow colors correspond to landmarks in lower panel). The bottom panel shows the distribution of mean emitted red values (represented by boxplots) across a sample of A, F1, and N individuals.

**Figure 7:**
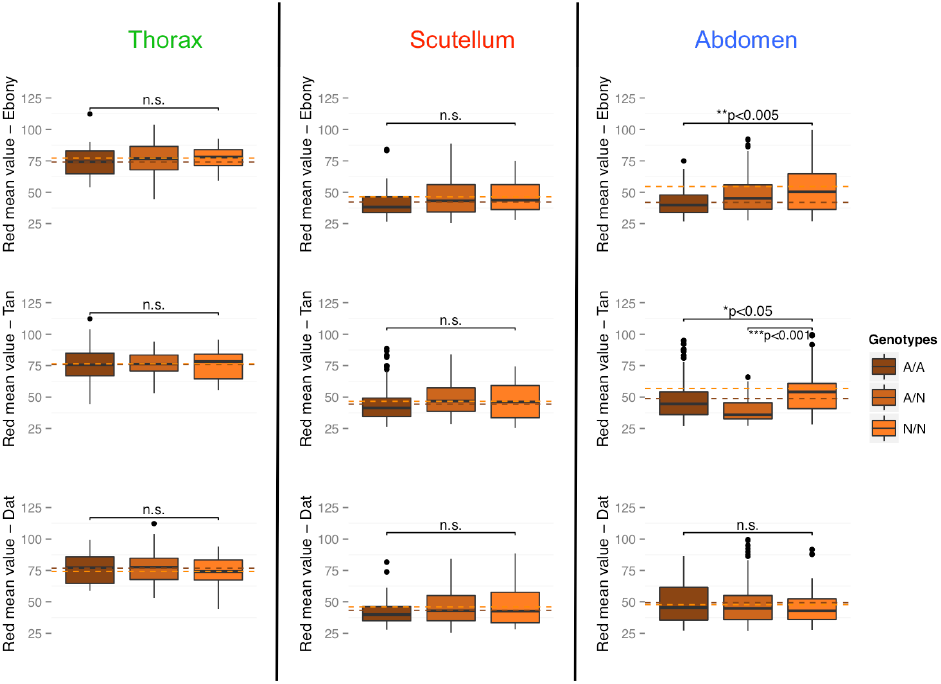
Boxplot distribution of mean red values for the three adult landmarks across genotype for the three pigmentation genes surveyed: Red mean values are partitioned into the three genotypic classes recovered in the F6 population (n=188). Dashed horizontal lines indicate the mean read value for A/A (brown) and N/N (orange) genotypes. Significant results from one-way ANOVA tests are indicated above each plot with asterisks and*p*-*value*, whereas non-significant results are labeled “n.s.”.

## Literature Cited

Ahmed-Braimah, Y. H., and B. F. McAllister, 2012 Rapid Evolution of Assortative Fertilization between Recently Allopatric Species of *Drosophila*. Int. J. of Evol. Bio. 2012: 1–9.

Arnoult, L., K. F. Y. Su, D. Manoel, C. Minervino, J. Magriña et al., 2013 Emergence and diversification of fly pigmentation through evolution of a gene regulatory module. Science 339: 1423–1426.

Bastide, H., A. Betancourt, V. Nolte, R. Tobler, P. Stöbe et al., 2013 A genome-wide, fine-scale map of natural pigmentation variation in Drosophila melanogaster. PLoS Genet. 9: e1003534.

Benson, G., 1999 Tandem repeats finder: a program to analyze DNA sequences. Nuc. Ac. Res. 27: 573–580.

Blight, W. C., and A. Romano, 1953 Notes on a breeding site of *Drosophila americana* near St-Louis, Missouri. Am. Nat. 87: 111–112.

Blumenstiel, J. P., 2014 Whole genome sequencing in *Drosophila virilis* identifies Polyphemus, a recently activated Tc1-like transposon with a possible role in hybrid dysgenesis. Mobile DNA 5: 6.

Brisson, J. A., D. C. De Toni, I. Duncan, and A. R. Templeton, 2005 Abdominal pigmentation variation in *Drosophila polymorpha:* Geographic in the trait, and underlying phylogeography. Evolution 59: 1046–1059.

Caletka, B. C., and B. F. McAllister, 2004 A genealogical view of chromosomal evolution and species delimitation in the *Drosophila virilis* species subgroup. Mol. Phyl. and Evol. 33: 664–670.

Clusella-Trullas, S., and J. S. Terblanche, 2011 Local adaptation for body color in *Drosophila americana:* commentary on Wittkopp et al. Heredity 106: 904–905.

Dai, F.-Y., L. Qiao, X.-L. Tong, C. Cao, P. Chen et al., 2010 Mutations of an arylalkylamine-N-acetyltransferase, Bm-iAANAT, are responsible for silkworm melanism mutant. J. Biol. Chem. 285: 19553–19560.

Fellowes, M. D. E., A. R. Kraaijeveld, and H. C. J. Godfray, 1999 The relative fitness of *Drosophila melanogaster* (Diptera, Drosophilidae) that have successfully defended themselves against the parasitoid *Asobara tabida* (Hymenoptera, Braconidae). J. of Evol. Bio. 12: 123–128.

Fonseca, N. A., R. Morales-Hojas, M. Reis, H. Rocha, C. P. Vieira et al., 2013 *Drosophila americana* as a model species for comparative studies on the molecular basis of phenotypic variation. Gen. Bio. and Evo. 5: 661–679.

Gloor, G, and W. R. Engels, 1992 Gloor: Single-fly DNA preps for PCR - Google Scholar. Dros. Info. Ser.

Haldane, J. B. S., 1937 The Effect of Variation of Fitness. Am. Nat. 71: 337–349.

Hazel, W., S. Ante, and B. Stringfellow, 1998 The evolution of environmentally-cued pupal colour in swallowtail butterflies: natural selection for pupation site and pupal colour. Ecol. Ent. 23: 41–44.

Hollocher, H., J. L. Hatcher, and E. G. Dyreson, 2000 Evolution of abdominal pigmentation differences across species in the *Drosophila dunni* subgroup. Evolution 54: 2046–2056.

Hsu, TC, 1952 Chromosomal Variation and Evolution in the Virilis Group of *Drosophila*. Univ. of Texas Publ. 5204: 35–72.

Jeong, S., M. Rebeiz, P. Andolfatto, T. Werner, J. True et al., 2008 The Evolution of Gene Regulation Underlies a Morphological Difference between Two *Drosophila* Sister Species. Cell 132: 783–793.

Livak, K. J., and T. D. Schmittgen, 2001 Analysis of Relative Gene Expression Data Using Real-Time Quantitative PCR and the 2-ΔΔCT Method. Methods 25: 402–408.

Matute, D. R., and A. Harris, 2013 The influence of abdominal pigmentation on desiccation and ultraviolet resistance in two species of *Drosophila*. Evolution 67: 2451–2460.

Morales-Hojas, R., M. Reis, C. P. Vieira, and J. Vieira, 2011 Resolving the phylogenetic relationships and evolutionary history of the *Drosophila virilis* group using multilocus data. Mol. Phyl. and Evol. 60: 249–258.

Muller, H. J., 1940 Bearings of the *Drosophila* Work on Systematics, pp. 185–268 in The New Systematics, Clarendon, Oxford.

Orr, H. A., 2001 The genetics of species differences. Trends in Eco. & Evol. 16: 343–350.

Orr, H. A., and J. A. Coyne, 1989 The genetics of postzygotic isolation in the *Drosophila virilis* group. Genetics 121: 527–537.

Orr, H. A., and J. A. Coyne, 1992 The Genetics of Adaptation - a Reassessment. Am. Nat. 140: 725–742.

Osanai-Futahashi, M., T. Ohde, J. Hirata, K. Uchino, R. Futahashi et al, 2012 A visible dominant marker for insect transgenesis. Nature Comm. 3.

Patterson, J. T., W. S. Stone, and A. B. Griffen, 1940 Evolution of the Virilis Group of *Drosophila*. Univ. of Texas Publ. 4032: 218–250.

Pool, J. E., and C. F. Aquadro, 2007 The genetic basis of adaptive pigmentation variation in *Drosophila melanogaster*. Mol. Eco. 16: 2844–2851.

Rozen, S, and H. Skaletsky, 2000 Primer3 on the WWW for general users and for biologist programmers. Methods Mol. Biol. 132: 365–386.

Sagga, N., and A. Civetta, 2011 Male-Female Interactions and the Evolution of Postmating Prezygotic Reproductive Isolation among Species of the Virilis Subgroup. Int. J. of Evol. Bio. 2011.

Spencer, WP, 1940 Subspecies, Hybrids and Speciation in *Drosophila Hydei* and *Drosophila Virilis*. Am. Nat. 74: 157–179.

Spicer, G. S., 1991 The genetic basis of a species-specific character in the *Drosophila virilis* species group. Genetics 128: 331–337.

Stalker, H. D., 1942 The inheritance of a subspecific character in the Virilis complex of *Drosophila*. Am. Nat. 76: 426–431.

Sweigart, A. L., 2010 The Genetics of Postmating, Prezygotic Reproductive Isolation Between *Drosophila virilis* and *D. americana*. Genetics 184: 401.

Takahashi, A., and T. Takano-Shimizu, 2011 Divergent enhancer haplotype of ebony on inversion In(3R)Payne associated with pigmentation variation in a tropical population of *Drosophila melanogaster*. Mol. Eco. 20: 4277–4287.

Takahashi, A., K. Takahashi, R. Ueda, and T. Takano-Shimizu, 2007 Natural variation of ebony gene controlling thoracic pigmentation in *Drosophila melanogaster*. Genetics 177: 1233–1237.

Telonis-Scott, M., A. A. Hoffmann, and C. M. Sgrò, 2011 The molecular genetics of clinal variation: a case study of ebony and thoracic trident pigmentation in *Drosophila melanogaster* from eastern Australia. Mol. Eco. 20: 2100–2110.

Terblanche, J. S., and T. E. Kleynhans, 2009 Phenotypic plasticity of desiccation resistance in Glossina puparia: are there ecotype constraints on acclimation responses? J. of Evol. Bio. 22: 1636–1648.

Trapnell, C., B. A. Williams, G. Pertea, A. Mortazavi, G. Kwan et al., 2010 Transcript assembly and quantification by RNA-Seq reveals unannotated transcripts and isoform switching during cell differentiation. Nature Biotech. 28: 511–U174.

Trapnell, C., L. Pachter, and S. L. Salzberg, 2009 TopHat: discovering splice junctions with RNA-Seq. Bioinformatics 25: 1105–1111.

True, J. R., 2003 Insect melanism: the molecules matter. Trends in Ecol. & Evo. 18: 640–647.

Werner, T., S. Koshikawa, T. M. Williams, and S. B. Carroll, 2010 Generation of a novel wing colour pattern by the Wingless morphogen. Nature 464: 1143–1148.

Williams, T. M., J. E. Selegue, T. Werner, N. Gompel, A. Kopp et al., 2008 The Regulation and Evolution of a Genetic Switch Controlling Sexually Dimorphic Traits in *Drosophila*. Cell 134: 610–623.

Wittkopp, P. J., K. Vaccaro, and S. B. Carroll, 2002 Evolution of yellow Gene Regulation and Pigmentation in *Drosophila*. Cur. Bio. 12: 1547–1556.

Wittkopp, P. J., S. B. Carroll, and A. Kopp, 2003a Evolution in black and white: genetic control of pigment patterns in *Drosophila*. Trends in Gen. 19: 495–504.

Wittkopp, P. J., B. L Williams, J. E. Selegue, and S. B. Carroll, 2003b *Drosophila* pigmentation evolution: Divergent genotypes underlying convergent phenotypes. Proc. to the Nat. Acad. of Sci.: 1–6.

Wittkopp, P. J., and P. Beldade, 2009 Development and evolution of insect pigmentation: Genetic mechanisms and the potential consequences of pleiotropy. Sem. in Cell & Dev. Bio. 20: 65–71.

Wittkopp, P. J., E. E. Stewart, L. L. Arnold, A. H. Neidert, B. K. Haerum et al, 2009 Intraspecific Polymorphism to Interspecific Divergence: Genetics of Pigmentation in *Drosophila*. Science 326: 540–544.

Wittkopp, P. J., G. Smith-Winberry, L. L. Arnold, E. M. Thompson, A. M. Cooley et al., 2010 Local adaptation for body color in *Drosophila americana*. Heredity 106: 592–602.

Zhan, S., Q. Guo, M. Li, M. Li, J. Li et al., 2010 Disruption of an N-acetyltransferase gene in the silkworm reveals a novel role in pigmentation. Development 137: 4083–4090.

